## Supplementary figures and images for "Change the direction: 3D optimal control simulation by directly tracking marker and ground reaction force data"

### Video Frontal View

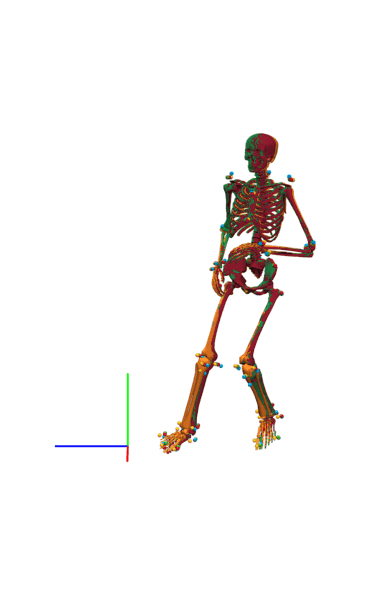

### Video Lateral View

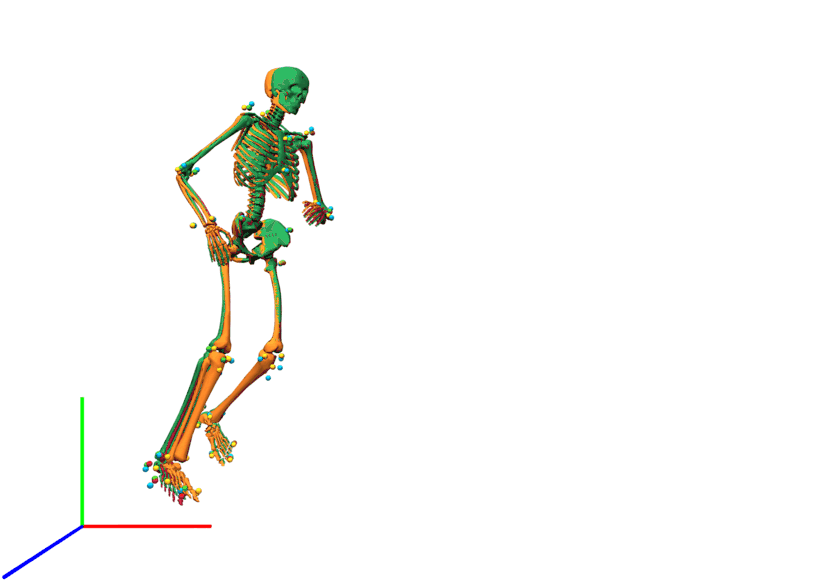
